# Doublecortin like kinase 1 (DCLK1) interacts with BRG1 chromatin remodeler and regulates neuronal activity-induced gene activation

**DOI:** 10.64898/2026.11.17.766006

**Authors:** Yi Luo, BongWoo Kim, Nasrin Haghighinajafabadi, Kenneth D Westover, Jiang Wu

**Affiliations:** Department of Physiology, The University of Texas Southwestern Medical Center, Dallas, TX75390, USA; Departments of Radiation Oncology and Biochemistry, The University of Texas Southwestern Medical Center, Dallas, TX 75390, USA

**Author notes:** These authors contributed equally (Yi Luo, BongWoo Kim). Correspondence: Jiang Wu, PhD Department of Physiology UT Southwestern Medical Center 5323 Harry Hines Blvd. Dallas TX 75390-9040.

## Abstract

SWI/SNF-like BAF ATP-dependent chromatin remodeling complexes play important roles in neural development and plasticity by regulating neural gene expression, especially in response to neuronal stimulations. Previously, we have shown that BRG1, the core subunit of BAF, undergoes neuronal activity-induced phosphorylation, which is important for the expression of a group of activity regulated genes (ARGs). Proteomic analyses identified Doublecortin like kinase 1 (DCLK1) as a novel BRG1 interacting protein. Neuronal depolarization induces nuclear enrichment of DCLK1 and increases interactions between BRG1 and DCLK1. We showed that the DCLK1 kinase domain and BRG1 C-terminal fragments are sufficient to mediate their interaction. Thus, DCLK1 is a candidate kinase responsible for BRG1 phosphorylation in response to neuronal activation. Using DCLK1 specific inhibitors and kinase inactive mutations, we showed that DCLK1, in a kinase activity dependent manner, promotes BRG1 phosphorylation and ARG activation. Thus, we identified a novel interacting kinase of BAF chromatin remodeling complex that regulates ARG expression. Because abnormal expression and mutations of both BAF complexes and DCLK1 are associated with neural disorders and cancer, our study provides molecular insights into their functions in these diseases.

## INTRODUCTION

The mammalian BAF complexes, also known as mSWI/SNF, are a group of ATP-dependent chromatin remodeling complexes consisting of core ATPase subunits BRG1/SMARCA4 or BRM/SMARCA2 along with 10-12 closely associated subunits. These remodelers utilize energy obtained through ATP hydrolysis to modify chromatin structures, thereby regulating transcription [1–3]. Mutations in BAF subunits are implicated as the genetic factors underlying Coffin-Siris syndrome, leading to severe neural developmental abnormalities [4–6]. Additionally, de novo functional mutations in genes encoding BAF subunits have been observed in patients with autism spectrum disorders, amyotrophic lateral sclerosis, and schizophrenia [7–10]. Notably, BRG1 serves as a crucial hub in the gene network associated with autism spectrum disorder [9]. Animal studies have also demonstrated significant roles of BAF subunits in neural development and plasticity [1, 2, 11].

BRG1 and BAF complexes exhibit a preference for binding to enhancers both during developmental processes and in cancer progression [12–14]. Although the biochemical and molecular mechanisms governing BAF function at enhancers are not fully characterized, it is known that BAF complexes can act as either activators or repressors depending on factors such as subunit composition, chromatin environment, and interactions with epigenetic regulators. Previous studies, including our own, have highlighted the context-dependent roles of neuronal BAF complexes in the regulation of expression of neuronal genes, especially their induction by neuronal activities [15–18]. However, the precise mechanisms through which BAF complexes exert specific functions in basal and activated neuronal conditions remain unclear.

Neuronal activities, eliciting Ca^2+^ influx via voltage-sensitive calcium channels and N-methyl-D-aspartate receptors, initiate multiple signaling pathways through intermediates, often kinases or phosphatases. These intermediates, including cAMP/PKA, Ras/MAPK, CaMK, and calcineurin, transmit signals to the nucleus, regulating transcription [19–22]. They phosphorylate or dephosphorylate numerous transcription factors and co-factors, acting as activity switches that govern ARG expression [22–24]. The phosphorylation of transcription regulators influences a dynamic protein interaction network, enabling neurons to swiftly and diversely respond, facilitating adaptation to the changing environment [25–28]. Recently, we identified BRG1-Ser1382 as a neuronal activity-induced phosphorylation site that was decreased by CaMKII inhibition. We showed that BRG1-pS1382 is important for BAF function in enhancer activation and neural plasticity in animal behavior. It remains largely unclear how this phosphorylation is regulated [29].

Doublecortin-like kinase 1 (DCLK1) contains two N-terminal doublecortin domains and a C-terminal CaMK-like kinase domain that function relatively independently [30, 31]. DCLK1 is highly enriched in brain and is critical in neural development. DCLK1 mutant mice displayed defects in neurite outgrowth and synaptic development [32]. Most adult non-neural tissues do not express detectable levels of DCLK1 protein. Interestingly, DCLK1 is found to be overexpressed in quiescent stem cells in several gastrointestinal cancers including pancreatic, esophageal, colon, and liver [33–35]. In other cancers, such as breast and kidney, DCLK1 is overexpressed and is important for cancer initiation, growth, stemness, the epithelial-mesenchymal transition, and metastasis [33, 36, 37]. The DCLK1 doublecortin domain binds to microtubules, and DCLK1 is found in both the cytoplasm and the nucleus. A recent study using DCLK1 specific chemical inhibitors and kinase inactive mutations revealed the connection between DCLK1 kinase activities and cancer growth by regulating RNA processing [38]. It is not clear whether the DCLK1 kinase domain is required for its role in neural development, and DCLK1 phosphorylation substrates remain largely unidentified.

To further determine how BAF complexes respond to neuronal activities and regulate ARG expression, we performed proteomic analyses of BAF complexes and interacting proteins in resting and depolarized neurons. We identified DCLK family of kinases as novel BAF interacting proteins that were affected by neuronal activation. Neuronal depolarization significantly increased nuclear levels of DCLK1 and the interactions between DCLK1 and BRG1. Using specific DCLK1 inhibitors and kinase inactive mutations, we demonstrated that DCLK1 promotes BRG1-S1382 phosphorylation and is important for ARG activation in a kinase activity-dependent manner. Thus, we identified a novel BAF interacting kinase that is induced by neuronal activities and regulates ARG activation.

## RESULTS

### Proteomic analyses identified DCLK1 as a potential neuronal activity-induced interacting protein of BRG1

Neuronal activities result in the activation of Ca^2+^ signaling. The subsequent activation of kinases such as calmodulin-activated CaMKs phosphorylate proteins in both nucleus and cytoplasm, affecting their interactomes and activities, which mediate neuronal physiological and transcription changes [19]. We performed proteomic analyses to determine how Ca^2+^ signaling could potentially affect BAF complex subunit composition, phosphorylation, and interacting proteins in response to neuronal activities. Using anti-BRG1/BRM antibody, we affinity-purified endogenous BAF complexes and interactomes from nuclear extracts of resting and depolarized cortical neuron cultures (Figure 1A). The mass spectroscopy analyses showed that the purifications yielded 272 proteins in the basal condition and 305 proteins in depolarized neuron samples [29]. These proteins include BRG1 (277 and 270 peptides, respectively) and most known BAF subunits and interacting proteins. Figure 1B shows representative BAF subunits and interacting proteins (see [29] for detail). Although there were no obvious subunit differences under the two conditions, several BAF interacting NuRD complex subunits were markedly reduced under the activated conditions. We have shown that the activity-induced BRG1 S1382 phosphorylation led to the reduced NuRD interaction and facilitates the departure of NuRD upon neuronal activation [29].

**Figure 1.**
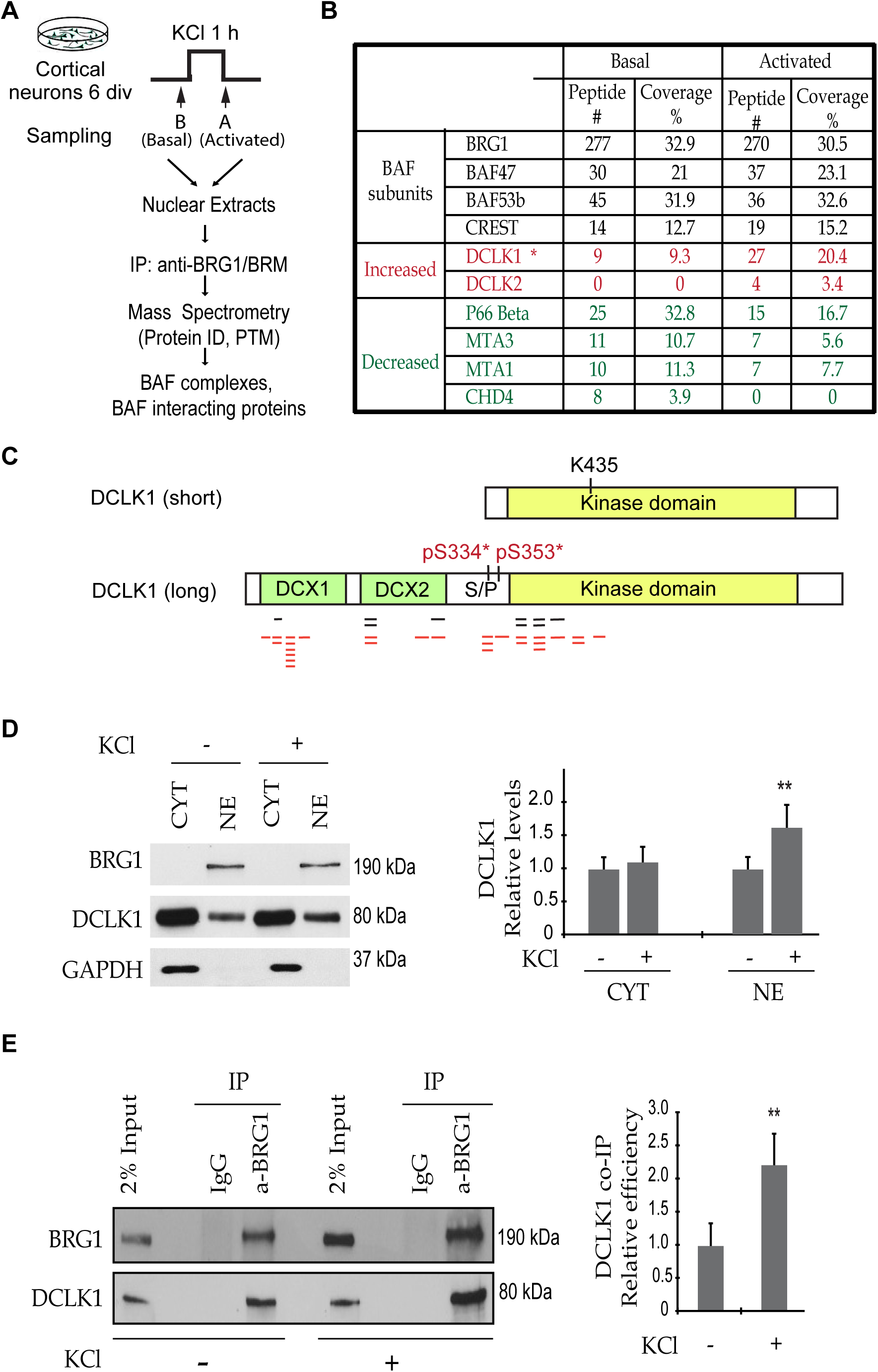
DCLK1 is a neuronal activity-induced interacting protein of BRG1. **A.** A diagram showing the purification and proteomic analyses of BAF complexes and interacting proteins from cortical neuron cultures under basal and depolarized conditions. **B.** Comparison of mass spec peptide numbers and coverages of representative BAF subunits and key dynamic interacting proteins. **C**. Domain structures of DCLK1 isoforms. Shown are the peptides identified from neurons under basal (in black) or depolarized (in red) conditions. The two phosphorylated serine sites that were identified from depolarized neurons are also shown. **D.** Western blot analyses of BRG1 and DCLK1 levels in the cytoplasmic and nuclear fractions of cortical neurons under basal or depolarized conditions. Quantifications are shown on the right. **E**. Western blot analyses of samples immunoprecipitated from nuclear extracts of neuron cultures in the absence or presence of KCl using antibodies against BRG1. Quantification of DCLK1 co-immunoprecipitation efficiency relative to BRG1 IP are shown on the right. Student’s t-test (n=3). **: p<0.01.

Interestingly, the most prominent BAF-interacting protein family that displayed potentially increased interaction with BRG1 was the doublecortin-like kinases (DCLK1 and DCLK2). We identified 9 peptides of DCLK1 under the basal condition and 27 peptides after KCl stimulation. DCLK2 peptides also increased from 0 to 4 (Figure 1B). DCLK1 has mainly two splicing isoforms with the short one lacking the microtubule binding doublecortin (DCX) domains. The DCLK1 peptides identified were distributed throughout the DCX domains and the kinase domain, suggesting that BRG1 likely interacts with the long form of DCLK1 in the nucleus (Figure 1C). Interestingly, in addition to BRG1 S1382 phosphorylation, two serine sites in the S/P rich hinge region of DCLK1 (S334 and S353) were identified as phosphorylated only in the complexes purified from the depolarized neurons (Figure 1C). These results suggest that DCLK1 is a neuronal activity-induced interacting protein of BRG1.

### Activity-induced increase of nuclear DCLK1 levels and DCLK1-BRG1 interactions

DCLK1 is largely a cytoplasmic protein and is also identified in the nucleus. To determine DCLK1 levels in different cell compartments and the effects of neuronal depolarization, we prepared cytoplasmic and nuclear extracts from cortical neuron cultures and performed western blot. The cytoplasmic and nuclear extracts were verified by the differential presence of cytoplasmic protein GAPDH and nuclear protein BRG1, respectively. We confirmed higher DCLK1 levels in the cytoplasmic fraction than in the nuclear fraction. The nuclear DCLK1 levels were increased in response to KCl treatment (Figure 1D). Using the nuclear extracts prepared from cortical neuron cultures under basal and depolarized conditions, we performed BRG1 immunoprecipitation (IP) coupled with western blot analyses. Endogenous nuclear DCLK1 was co-IPed with BRG1. Normalized with IPed BRG1 levels, the co-IP efficiency of DCLK1 displayed a 2-fold increase after KCl treatment (Figure 1E). Thus, the increased DCLK1-BRG1 interaction could be contributed by both increased nuclear DCLK1 levels and increased DCLK1-BRG1 affinities in response to neuronal activation.

### Domains involved in the DCLK1-BRG1 interaction

BRG1 contains an N terminal specific region, the HSA domain that is important for core subunits assembly, the ATPase domain, the SnAc domain that is important for histone anchor, and the bromo domain that may target BAF complex to specific chromatin locations (Figure 2A). The activity-induced phosphorylation occurs on S1382, which is right downstream of the SnAc domain. Notably, the C-terminal region downstream of the SnAc domain was not present in the Cryo-EM structure of the BAF complex [39, 40], which suggests that this region is likely a flexible region mediating the interactions with other proteins. To determine which parts of BRG1 protein interact with DCLK1, we generated constructs expressing full length and different truncated proteins in HeLa cells. IP-western blot experiments were then performed (Figure 2A, 2B). The BRG1 specific antibody (G7 clone, Santa Cruz) recognizes the specific N-terminal domain of BRG1. We used HA tag to detect the C-terminal of BRG1. Whereas the N-terminal domain (BRG1 1-399) did not co-IP with DCLK1, other BRG1 fragments including the C-terminal fragments did. In a reciprocal experiment, we performed immunoprecipitation using either a BRG1 antibody or anti-HA antibodies. In both conditions, DCLK1 could interact with full length BRG1 and BRG1 C-terminus (Figure 2C).

**Figure 2.**
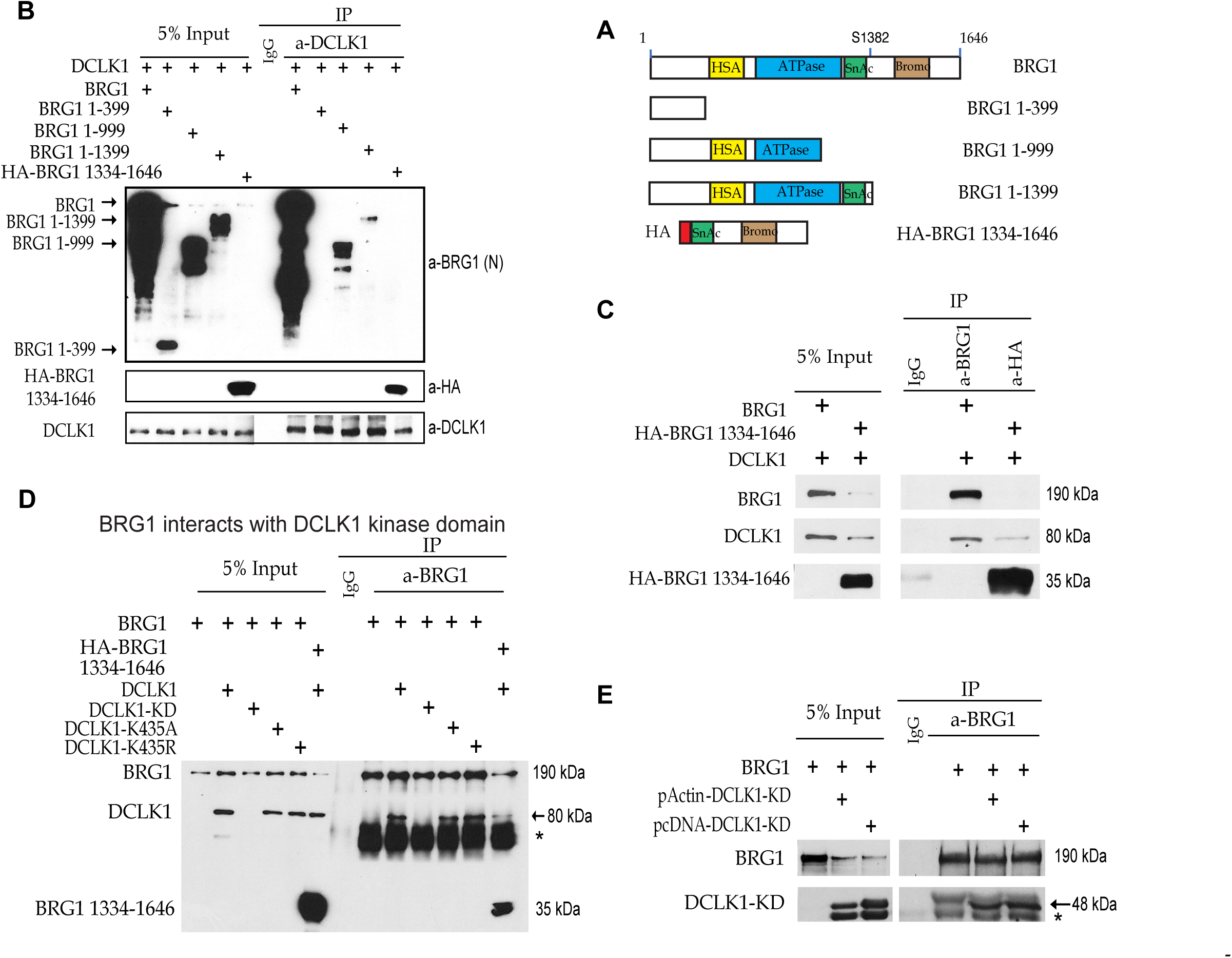
Domains involved in the DCLK1-BRG1 interaction. **A**. A diagram illustrating domain structures of full length BRG1 and BRG1 truncated fragments used in the interaction study. **B**. IP-western blot analyses of BRG1 fragments that co-IPed with full length DCLK1 proteins. **C**. IP-western blot analyses of DCLK1 that co-IPed with full length and C-terminal BRG1. **D**. IP-western blot analyses of wildtype and kinase inactive form of DCLK1 that co-IPed with full length and C-terminal BRG1. **E**. IP-western blot analyses showed interactions between DCLK1 kinase domain (KD) and BRG1. *, IgG.

BRG1 C-terminus includes S1382, suggesting that DCLK1 is a candidate kinase for S1382. To determine whether DCLK1 kinase domain and kinase activities are involved in the interaction with BRG1, we generated a DCLK1 kinase domain fragment (DCLK1-KD) and two kinase inactive mutations (K435A and K435R[32]) (Figure 2D, 2E). Wildtype as well as kinase inactive DCLK1 co-IPed with both full length and the C-terminus of BRG1 (Figure 2D). The kinase domain of DCLK1 was also sufficient to interact with BRG1 (Figure 2E). Thus, DCLK1 could interact with BRG1 through the kinase domain of DCLK1 and the C-terminus of BRG1.

### DCLK1 promotes BRG1-S1382 phosphorylation

We discovered that BRG1 S1382 is phosphorylated in response to various neuronal activities [29]. We also identified CaMKII as a candidate kinase. The neuronal activity-induced BRG1-DCLK1 interaction and the presence of a CaMK like kinase domain in DCLK1 suggest that DCLK1 could potentially also phosphorylate BRG1-S1382. We first co-expressed BRG1 and different DCLK1 forms in HeLa cells and compared BRG1 migration distance in SDS-PAGE (Figure 3A). It appears that BRG1 migration was slower than controls when co-expressed with either full length DCLK1 or DCLK1 kinase domain (DCLK1-KD). The co-expression of DCLK1 kinase inactive mutants did not affect BRG1 migration. β-catenin was used as a control for protein migration in the same gel (Figure 3A). This DCLK1-caused BRG1 slow migration was lost after calf intestinal alkaline phosphatase (CIP) treatment of the cell lysates (Figure 3B), which suggest that the slower BRG1 migration was caused by BRG1 phosphorylation.

**Figure 3.**
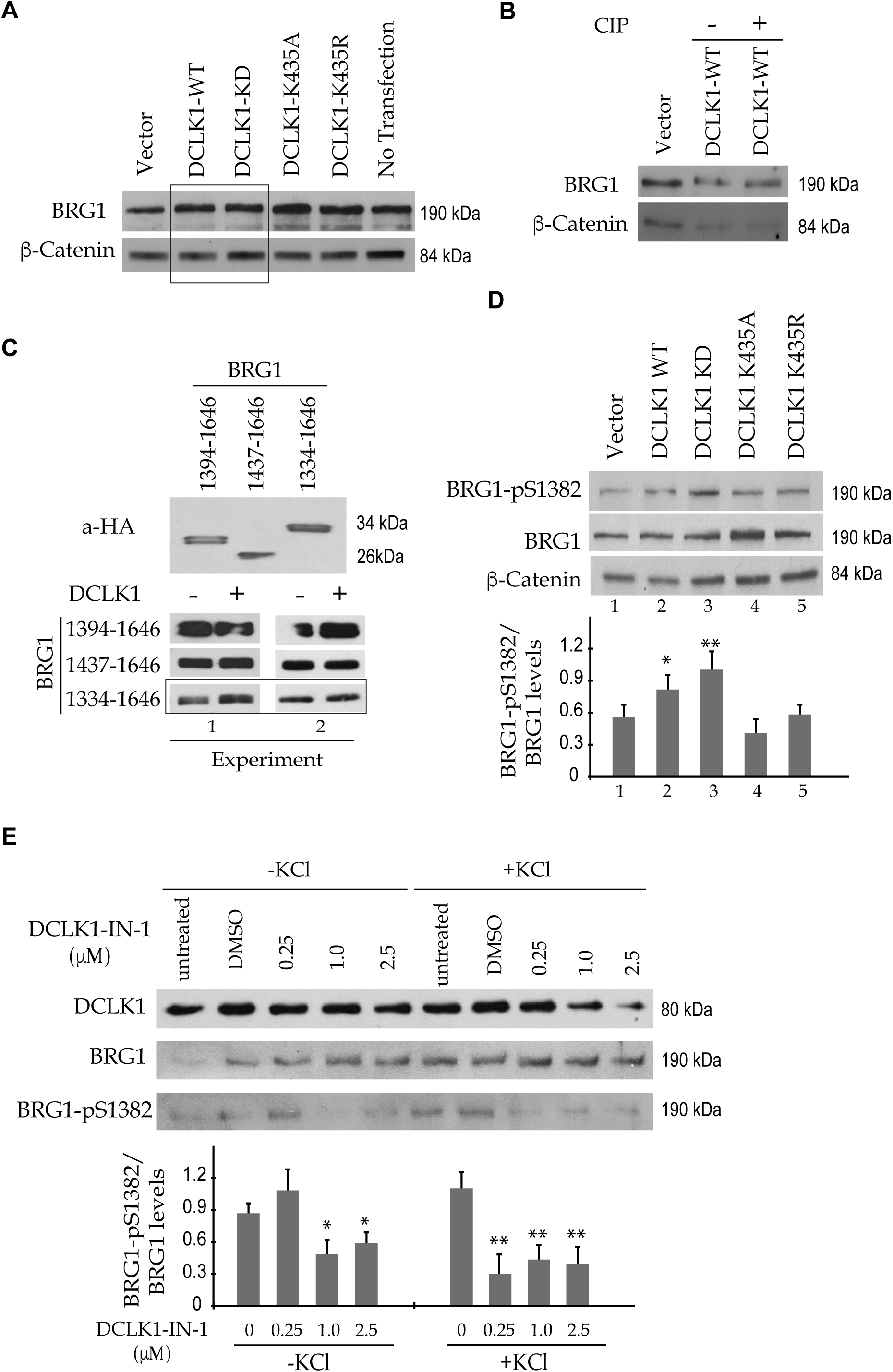
DCLK1 promotes BRG1-S1382 phosphorylation. **A.** Western blot analyses show slower BRG1 migration when co-expressed with full length wildtype DCLK1 or the DCLK1 kinase domain (DCLK1-KD) (boxed), but not with kinase inactive DCLK1. β-Catenin was used as a gel running control. **B**. Western blot analyses showing the migration change of BRG1 co-expressed with DCLK1 with or without CIP treatment. **C**. Western blot analyses showing the expression of BRG1 C-terminal fragments in HeLa cells (top panel) and migration comparisons when co-expressed with DCLK1. Note the change of BRG1 1334-1646 fragment migration upon DCLK1 co-expression (boxed). **D**. Western blot showing BRG1-pS1382 levels in synchronized HeLa cells at G0/G1 phase co-expressing BRG1 and different DCLK1 forms. **E**. Western blot showing endogenous BRG1-p1382 levels in cortical neuron cultures under basal and KCl treatment (50 mM, 1 hour). DCLK1-IN-1 at indicated concentration or DMSO solvent controls were added before KCl treatment. Quantifications of BRG1-pS1382/BRG1 ratio in **D** and **E** are shown below the blots. Student’s t-test compared to the control conditions (n=3). *: p<0.05. **: p<0.01.

To determine whether the phosphorylation of BRG1 occurs on S1382, we first co-expressed DCLK1 with different lengths of BRG1 C-terminal fragments. Indeed, only BRG1-1334-1646 which included S1382, but not the other two smaller fragments that excluded S1382, migrated slower when co-expressed with DCLK1 (Figure 3C). To specifically examine whether DCLK1 could phosphorylate BRG1-S1382, we performed BRG1 western blot using the BRG1-pS1382 phospho-specific antibodies [29]. BRG1 S1382 phosphorylation was increased when BRG1 was co-expressed with either full length wild type DCLK1 or DCLK1 kinase domain. The co-expression of DCLK1 kinase inactive mutants did not affect BRG1-pS1382 levels (Figure 3D). In primary cultured neurons, in which depolarization induced S1382 phosphorylation of endogenous BRG1, pre-treatment with a DCLK1-specific inhibitor (DCLK1-IN-1) [41] reduced both DCLK1 and BRG1-pS1382 levels (Figure 3E). These results suggest that DCLK1 could promote BRG1-S1382 phosphorylation, especially in response to neuronal activities.

### Neuronal activity dependent ARG transcription was impaired by DCLK1 inhibitors

We have shown that BRG1 and its phosphorylation promote neuronal activity-induced gene activation [29]. To determine whether DCLK1 are important for the activation of ARGs, we used DCLK1 inhibitors. Using a previously identified LRRK2-IN-1 that also targets DCLK1[42], we examined the effects of inhibiting DCLK1 in ARG activation. Treating cultured cortical neurons with KCl led to increased expression of prototypical immediate early genes (IEGs) and ARGs including *c-Fos* and *Bdnf*. LRRK2-IN-1 treatment resulted in impaired *c-Fos* and *Bdnf* induction in a dosage dependent manner (Figure 4A). Similar to a previous report [42], LRRK2-IN-1 led to the decreases DCLK1 protein levels in a dosage-dependent manner as shown in western blots (Figure 4B). Recently, DCLK1-IN-1, a more specific DCLK1 inhibitor has been developed [41], which has an effective range of 0.25 to 2.5 μM for DCLK1 inhibition (Figure 3E). Interestingly, treating cortical neuron cultures with DCLK1-IN-1 at a suboptimal dose of 0.25 μM led to increased activation of *c-Fos* and *Arc*, but at 1 and 2.5 μM, where DCLK1 protein was reduced to lower levels (Figure 3E), ARG activation was impaired. Thus, DCLK1 has dose-dependent roles in ARG activation.

**Figure 4.**
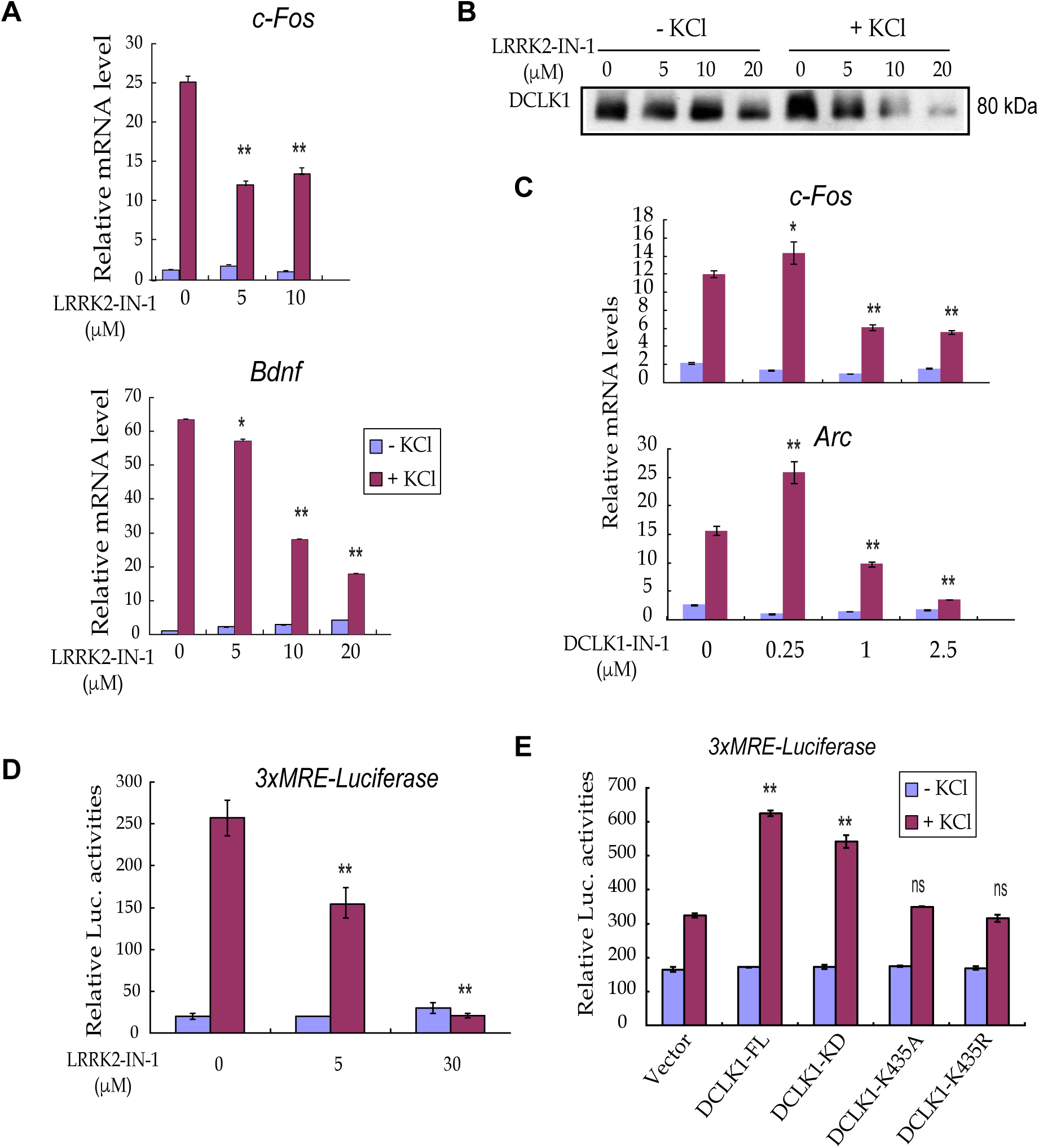
DCLK1 regulates neuronal activity dependent ARG transcription. **A.** Relative mRNA levels of *c-Fos* and *Bdnf* in cultured neurons with or without KCl treatment (50 mM, 4 hr) as shown by RT-qPCR. DCLK1 inhibitor LRRK2-IN-1 at indicated concentrations or solvent control were added before KCl treatment. **B**. Western blot analyses showing endogenous levels of DCLK1 in neurons treated with LRRK2-IN-1 and KCl as described in **A**. **C.** Relative mRNA levels of *c-Fos* and *Arc* in cultured neurons with or without KCl treatment (50 mM, 4 hr) as shown by RT-qPCR. DCLK1 inhibitor DCLK1-IN-1 at indicated concentrations or solvent control were added before KCl treatment. **D.** Relative Luciferase activities in neurons transfected with 3xMRE-Luciferase reporter genes with or without KCl treatment (30 mM, 24 hr). LRRK2-IN-1 at indicated concentrations or solvent control were added before KCl treatment. **E**. Relative Luciferase activities in neurons transfected with 3xMRE-Luciferase reporter genes and different DCLK1 forms with or without KCl treatment (30 mM, 24 hr). Student’s t-test compared to DMSO treated conditions (n=3). *: p<0.05. **: p<0.01.

### Function of DCLK1 in ARG activation requires its kinase activity

Previously we have shown that BRG1 functions together with a neuronal activity-regulated transcriptional factor MEF2C and is required for MEF2C-mediated target gene activation [17]. We transfected a reporter system with luciferase expression driven by 3xMEF2C response element (3xMRE) into cultured cortical neurons. We showed that the activity-induced luciferase expression in cultured cortical neurons treated with KCl was impaired by increased dose of LRRK2-IN-1 (Figure 4D). This result is consistent with our observations with the endogenous ARGs. To determine whether DCLK1 is sufficient to promote ARG activation, we overexpressed full length DCLK1 in cultured neurons. When treated with KCl, the induction levels of reporter luciferase levels were significantly increased compared to neurons transfected with empty vectors. Importantly, overexpressing the kinase domain of DCLK1 also increased 3xMRE luciferase expression. On the other hand, the two kinase inactive mutants of DCLK1 failed to increase the KCl induced luciferase expression (Figure 4E). These results suggest that function of DCLK1 in promoting ARG induction requires its kinase activity.

## MATERIALS AND METHODS

### Mice

Wildtype C57BL/6 mice were housed at the UT Southwestern Medical Center Animal Facility, and all procedures followed IACUC-approved protocols. In all animal experiments, both male and female embryos were used, and there were no significant differences found between them.

### Cortical neuron culture, inhibitor treatment, and transfection

E16.5 cortical cells from wildtype C57BL/6 mice were cultured as previously described [16, 17]. Dissociated cortical neurons were plated on poly-L-ornithine- and fibronectin-coated wells. Culture media contained neurobasal plus with B27 plus supplement (Gibco). Cortical cultures were transfected with plasmids on 4 days *in vitro* (div) and analyzed at 6 div. For depolarization, the cultures were pretreated with TTX (1 µM) and APV (100 µM) to inhibit spontaneous activation on 5 div and 50 mM KCl was added to the cultures for 1 to 6 h on 6 div as described [43]. For 3xMRE reporter assays, 30 mM KCl were used for 24 hr treatment for better neuron survival and sufficient luciferase production [17]. For inhibitor treatment, LRRK2-IN-1, DCLK1-IN-1, or DMSO (as a control) with indicated concentrations was added on 5 div before KCl treatment on 6 div. Cortical cultures were transfected with reporter and DCLK1 constructs with Lipofectamine (Thermo Fisher) on 5 div and analyzed at 6 div before or after 30 mM KCl treatment. Luciferase reporter assays were performed using the Dual-Luciferase Assay kit (Promega).

### Affinity purification of BAF complexes and mass spectrometry analysis

The proteomic analyses of BAF complexes were performed as described in [29]. Briefly, nuclear extracts prepared from 6-div mouse cortical neurons with or without KCl (50 mM) treatment for 1 h were used for affinity purification with the anti-BRG1 J1 antibody or rabbit IgG control cross-linked to GE FastFlow Sepharose Protein A beads. The eluted BAF complexes and associated proteins from gel slices were subjected to LC-MS/MS, which was performed by PTM Biolabs (Chicago, US). The resulting MS/MS data were processed using the Mascot search engine (version 2.3). Tandem mass spectra were searched against SwissProt database concatenated with reverse decoy database. Trypsin/P was specified as cleavage enzyme allowing up to 3 missing cleavages. Mass error was set to 10 ppm for precursor ions and 0.02 Da for fragment ions. Phosphorylations of Ser, Thr, and Tyr were specified as variable modifications. An FDR of less than 1% was used to filter the identified peptides. All the other parameters in Mascot were set to default values.

### HeLa cell culture, plasmid and transfection

HeLa cells (ATCC CCL-2) were cultured in DMEM (Invitrogen) supplemented with 10% FBS and 2 mM L-glutamine. HeLa cell G0/G1 synchronization was achieved by serum starvation for 24 hours. The MEF2 reporter construct MRE-Luciferase contains three MRE sequences upstream of the luciferase reporter [17]. pSin4-EF2-IRES-Puro was used to clone BRG1 [44] and BRG1 fragments for expression of BRG1 in cultured cells. DCLK1 and mutant fragments were cloned in the expression vectors pActin or pcDNA3.1. PolyJet (Signagen) was used for plasmid transfection of cultured HeLa cells.

### RT-PCR and q-PCR

RNA from cells was extracted with TRIZOL (Invitrogen). cDNAs were synthesized by reverse transcription using Iscript (Bio-Rad), followed by PCR or quantitative PCR analysis. A Bio-Rad real-time PCR system (C1000 Thermal Cycler) was used for quantitative PCR. Levels of *GAPDH* mRNA were used to normalize input RNA. Graphics shown are representative of experiments performed in triplicate. The experiments were repeated at least three times. Standard errors were calculated according to a previously described method [44]. The sequences of all the primers are *c-Fos* (5’ ATCCTTGGAGCCAGTCAAGA3’, and 5’ ATGATGCCGGAAACAAGAAG3’), *Arc* (5’ GTGAAGACAAGCCAGCATGA3’ and 5’ CCAAGAGGACCAAGGGTACA3’), and *Bdnf* (5’CAGTACAGTGGTTCTACAATC3’, and 5’ CAATTCACAATTAAAGCAGC3’).

### Immunoprecipitation and western blot

Cortical cultures expressing endogenous BRG1 and DCLK1 or HeLa cells expressing exogenous BRG1 and DCLK1 fragments were lysed with Buffer A (25 mM Tris, pH 7.5, 25 mM KCl, 5 mM MgCl_2_, 10% glycerol, 0.1% NP-40, with protease inhibitor freshly added). Nuclear extracts were prepared in RIPA buffer (150 mM NaCl, 1.0% NP-40, 0.5% sodium deoxycholate, 0.1% SDS, and 50 mM Tris, pH 8.0) with rotation at 4 °C for 1 h. After centrifugation, rabbit polyclonal antibodies against BRG1/BRM (J1) [45], DCLK1 (Abcam ab31704), or HA tag (Abcam, ab9110) were added to pre-cleared nuclear extracts, and samples were incubated at 4 °C overnight. Samples were incubated with protein A beads (GE Healthcare) for 1 h; beads were washed with RIPA buffer four times. Precipitated proteins were eluted by boiling in 2X Sample Buffer (Bio-Rad) before SDS-PAGE and western blot analysis. For immunoblotting, cell lysates or immunoprecipitation were separated on SDS-PAGE gels. Antibodies used were against BRG1 (G7, Santa Cruz Biotechnology), BRG1-pS1382 [29], β-catenin (Abcam ab6302), GAPDH (Invitrogen PA1-988) and DCLK1 (Abcam ab31704). HRP-conjugated secondary antibodies were purchased from Jackson Immunology. Quick CIP kit (NEB, M0525) was used for treating protein lysates to dephosphorylate proteins.

### Statistical analysis

Unless otherwise specified, at least three independent experiments were performed and each condition was analyzed in triplicate. Bar graphs shown are representative experiments. Data are expressed as means ± s.d. Statistical analysis was performed by either analysis of variance with ANOVA post hoc t-test for multiple comparisons or a two-tailed unpaired Student’s t-test. A p value of <0.05 was considered significant.

## DISCUSSION

In this study, we identified DCLK family of kinases as novel interacting proteins of the SWI/SNF like BAF complexes. Focusing on DCLK1, we show that the interaction was significantly enhanced by neuronal activation, and DCLK1 plays an important role in regulating neuronal activity-induced gene expression. Neuronal activities may regulate DCLK1-BAF interaction in several levels. We showed that neuronal depolarization led to increased DCLK1 protein levels in the nucleus. In addition, we identified 2 serine phosphorylation sites in DCLK1 that are induced by neuronal activities and are in the hinge region, which has been shown to regulate DCLK1 activities [46]. It remains to be tested if the modifications could increase DCLK1-BAF interactions and/or DCLK1 kinase activities. Notably, a similar proteomic analysis of BAF interaction proteins in neurons identified DCLK1/2 as BAF-interacting proteins that could be induced by neuronal depolarizations [47]. It is possible but not tested whether DCLK2 functions similarly to DCLK1. Both DCLK1 inhibitors used in the study could not distinguish DCLK1 and DCLK2.

BAF subunits undergo phosphorylation by many kinases in response to several stimulations. During mitosis, BRG1 phosphorylation leads to its dissociation from condensed chromatin and subsequent inactivation [48, 49]. It has been shown that Casein kinase 2 phosphorylates BRG1 at multiple sites during mitosis, with these phosphorylation events playing a crucial role in myoblast proliferation and survival [26, 50]. During myogenesis, BRG1 phosphorylation by PKCβ1 and dephosphorylation by calcineurin actively regulate gene expression [51]. In Drosophila, BRM phosphorylation by cyclin-dependent kinases is essential for cell proliferation and the development of wing epithelium[52]. Beyond cell-cycle progression and development, BRG1 is phosphorylated in response to DNA damage signaling by the ATM kinase, leading to the binding of the ATPase to γ-H2AX-containing nucleosomes [25]. Additionally, BAF60 undergoes phosphorylation by p38 during muscle development [53]. PHF10/BAF45a and BAF155 can also be phosphorylated in various cell types [54, 55]. We have shown that calcium signaling pathways mediated by CaMKII regulate BRG1 phosphorylation at S1382 in postmitotic neurons [29]. The activity-induced BRG1-DCLK1 interaction suggest that DCLK1 could be an additional kinase that phosphorylates BRG1-S1382. We showed that the DCLK1 kinase domain and the C-terminus of BRG1 were sufficient for their interaction (Figure 2). Co-expression of DCLK1 or its kinase domain with BRG1 could increase BRG1-pS1382 (Figure 3). Inhibition of DCLK1 in neurons significantly reduced BRG1-pS1382. Alternatively, CaMKII could regulate BRG1-pS1382 through regulating DCLK1 activities. More study will be needed to reveal the functional and biochemical interactions between BRG1 and these activity-regulated kinases. In addition to BRG1-S1382, DCLK1/2 may be able to phosphorylate other BAF subunits and additional chromatin regulators. DCLK1-IN-1 inhibitor was reported to reduce the activity-induced phosphorylation of several BAF subunits including BRG1 [56].

DCLK1 expression is highly enriched in the brain during normal development. It plays critical roles in neural development and function. Studies linked its microtubule binding doublecortin domains and function in neuronal morphogenesis and migration [30]. DCLK1 has been shown to regulate α-synuclein levels and potentially neurodegenerative disease [57]. DCLK1 knockdown and knockout neurons display dendrite and synapse development defects, and the kinase domains are involved in synapse maturation [32]. DCLK1 genetic variants in humans are associated with mental disorders [58]. Our study identified a novel function of DCLK1 in neuronal transcription regulation (Figure 4). Neuronal activity could induce a nuclear enrichment of DCLK1 and its interactions with BAF complexes. The small molecule DCLK1 inhibitors inhibited the activity-induced expression of both 3xMRE-Luc reporter and endogenous ARGs in a dosage-dependent manner. Overexpression of DCLK1 and its kinase domain, but not overexpression of the kinase-inactive mutants, enhanced the response of 3xMRE-Luc to neuronal activation. These data suggest a function of DCLK1 in activity-dependent gene regulation, potentially through its interaction with and regulating the BAF chromatin remodeling complex. Both BAF complexes and DCLK1 are important players in various cancers, it remains to be tested whether BAF-DCLK1 interactions occur in the cancer contexts.

In summary, we identified DCLK1 as a novel neuronal activity-induced BRG1 interacting kinase. We demonstrated that DCLK1 promotes BRG1 phosphorylation. We revealed a novel function of DCLK1 in regulating neuronal activity-induced gene activation, which is dependent on its kinase activities. Our study provides insights into the function and regulation of BAF complexes and DCLK1, both of which are important players in neural development and neural disorders.

## Acknowledgements

We are grateful to Dr. Nathanael S Gray (Stanford University) for providing us DCLK1-IN-1. This work was supported by grants from the NIH (R01NS137584 and R01NS096068 to J.W.) and CPRIT (RP220235 to J.W.).

## Contributions

J.W. and Y.L. designed the experiments. Y.L., B.K., and N.H. performed the experiments and collected the data. Y.L., B.K. and J.W. analyzed the results. K.D.W provided reagents and discussion. J.W. wrote the manuscript with the help from all authors.

